# Laser nanobubbles induce immunogenic cell death in breast cancer

**DOI:** 10.1101/2020.09.04.283846

**Authors:** Hieu T.M. Nguyen, Nitesh Katta, Jessica A. Widman, Eri Takematsu, Xu Feng, Susana H. Torres, Tania Betancourt, Aaron B. Baker, Laura J. Suggs, Thomas E. Milner, James W. Tunnell

## Abstract

Recent advances in immunotherapy have highlighted a need for therapeutics that initiate immunogenic cell death in tumors to stimulate the body’s immune response to cancer. This study examines whether laser-generated bubbles surrounding nanoparticles (“nanobubbles”) induce an immunogenic response for cancer treatment. A single nanosecond laser pulse at 1064 nm generates micron-sized bubbles surrounding gold nanorods in the cytoplasm of breast cancer cells. Cell death occurred in cells treated with nanorods and irradiated but not in cells with irradiation treatment alone. Cells treated with nanorods and irradiation had increased damage-associated molecular patterns (DAMPs), including increased expression of chaperone proteins human high mobility group box 1 (HMGB1), adenosine triphosphate (ATP), and heat shock protein 70 (HSP70). This enhanced expression of DAMPs led to the activation of dendritic cells. Overall, this treatment approach is a rapid and highly specific method to eradicate tumor cells with simultaneous immunogenic cell death signaling, showing potential as a combination strategy for immunotherapy.

---

Immunotherapy has become the primary treatment for several advanced, metastatic cancers including melanoma, lung cancer, and head and neck cancers.^1–4^ Despite the success of immunotherapy, the proportion of patients that do not respond or have incomplete responses to the therapy are high for many types of cancer.^3,5,6^ The reason for these low response rates is believed to be that tumors produce immunosuppressive factors that prevent immune recognition and tumor cell death.

A key strategy to enhance immunotherapy is to elicit immunogenic cell death in tumor cells. Immunogenic cell death results in both antigenicity (release of tumor-specific antigen) and adjuvanticity (release of molecular signaling that stimulates immune responses). Adjuvant signaling by the secretion of damage-associated molecular patterns (DAMPs) has been shown to activate dendritic cells to acquire tumor-specific antigens that mount adaptive T cell responses specific to tumor cells.^7–9^

Targeted hyperthermia (locally controlled tumor radiative heating with laser ^10–12^) has emerged as a promising therapeutic approach that elicits immunogenic tumor cell death.^13–17^ Most recently, the synergy between hyperthermia therapy and immunotherapy (immune checkpoint and adoptive T cell therapy) was demonstrated in pre-clinical models illustrating that, when combined with hyperthermia, tumor burden was minimized over any monotherapy.^18–20^

Laser nanobubbles (bubbles generated around nanoparticles from irradiation with nanosecond pulsed laser radiation) offer an alternative method to trigger cell death via physical disruption of cell membranes. This mechanism leads to a necrotic cell death fate^21^ with the potential to elicit more inflammatory, pro-immunogenic signaling. The secretion of immunogenic markers into the extracellular environment by laser nanobubbles occurs in binary events and without strong dependence on dosimetry. Therefore, they eliminate the need for a dosimetry monitoring system during laser treatment. Moreover, laser nanobubbles can trigger cell death after one pulse of laser irradiation,^21–24^ while targeted hyperthermia typically requires a few minutes to deliver the optimal temperature.^25^ The rapid therapeutic creation of laser nanobubbles may facilitate the treatment of large tumors.

In this study, we demonstrate immunogenic cell death from laser nanobubbles for the first time. Following a single nanosecond laser pulse irradiation, rapid breast cancer cell death occurred due to membrane disruption. Moreover, this effect was highly specific, causing membrane disruption only in cells with gold nanorods (AuNRs), while neighboring cells without AuNRs were left intact. We also observed bubble formation in cells, confirming the AuNRs-laser interaction is transient and discrete. We determined that extracellular release of DAMPs, including chaperone proteins, human high mobility group box 1 (HMGB1), adenosine triphosphate (ATP), and heat shock protein 70 (HSP70) were enhanced in the laser treatment group. With the presence of DAMPs secreted from laser irradiation, dendritic cell activation increased. Overall, we demonstrated that nanosecond pulsed laser irradiation provided a fast and highly specific therapy to eradicate tumor cells and elicit immunogenic cell death, highlighting the potential of this approach as a candidate combination strategy for immunotherapy.

We used AuNRs coated with (11-Mercaptoundecyl)-N,N,N-trimethylammonium bromide (Mutab), a quaternary ammonium compound that is capable of driving cellular uptake due to their positive zeta potential.^26,27^ The AuNRs were 38 nm long and 10 nm wide (aspect ratio of 4:1) and had a peak surface plasmon resonance (SPR) at 788 nm (Figure 1a). Laser irradiation at 1064 nm only results in 25% absorption of the SPR peak at 788 nm. While the longitudinal SPR peak of AuNRs can be tuned to match the laser irradiation by increasing the AuNRs aspect ratio, cellular uptake significantly reduces with increasing AuNR length.^28–30^ This off-resonance absorption can be compensated by radiation with higher fluence while maintaining a significant margin between triggering cell death with and without AuNRs^23^. The metabolic activities of human and murine breast cancer cells (MDA-MB-231 and 4T1 cells) were characterized by the tetrazolium compound [3-(4,5-dimethylthiazol-2-yl)-5-(3-carboxymethoxyphenyl)-2-(4-sulfophenyl)-2H-tetrazolium (MTS) assay. The metabolic activities of cells incubated with AuNRs within 3, 6, 12, and 24 hours were similar to that of cells without AuNRs (Figure 1b). Hence, AuNRs have minimal cytotoxicity on MDA-MB-21 and 4T1 cells. Two-photon microscopy images of AuNRs internalization in MDA-MB-231 and 4T1 cells shows the uptake of AuNRs for 6 hours (Figure 1c). The cytoplasm of live cells stained with calcein-AM appears in green, while AuNRs clusters photo-luminesce over a broad spectrum and appear in yellow as a result of the overlapping of red and green channels. The internalized AuNRs cluster in various sizes and appear randomly distributed in the cytoplasm.

**Figure 1.**
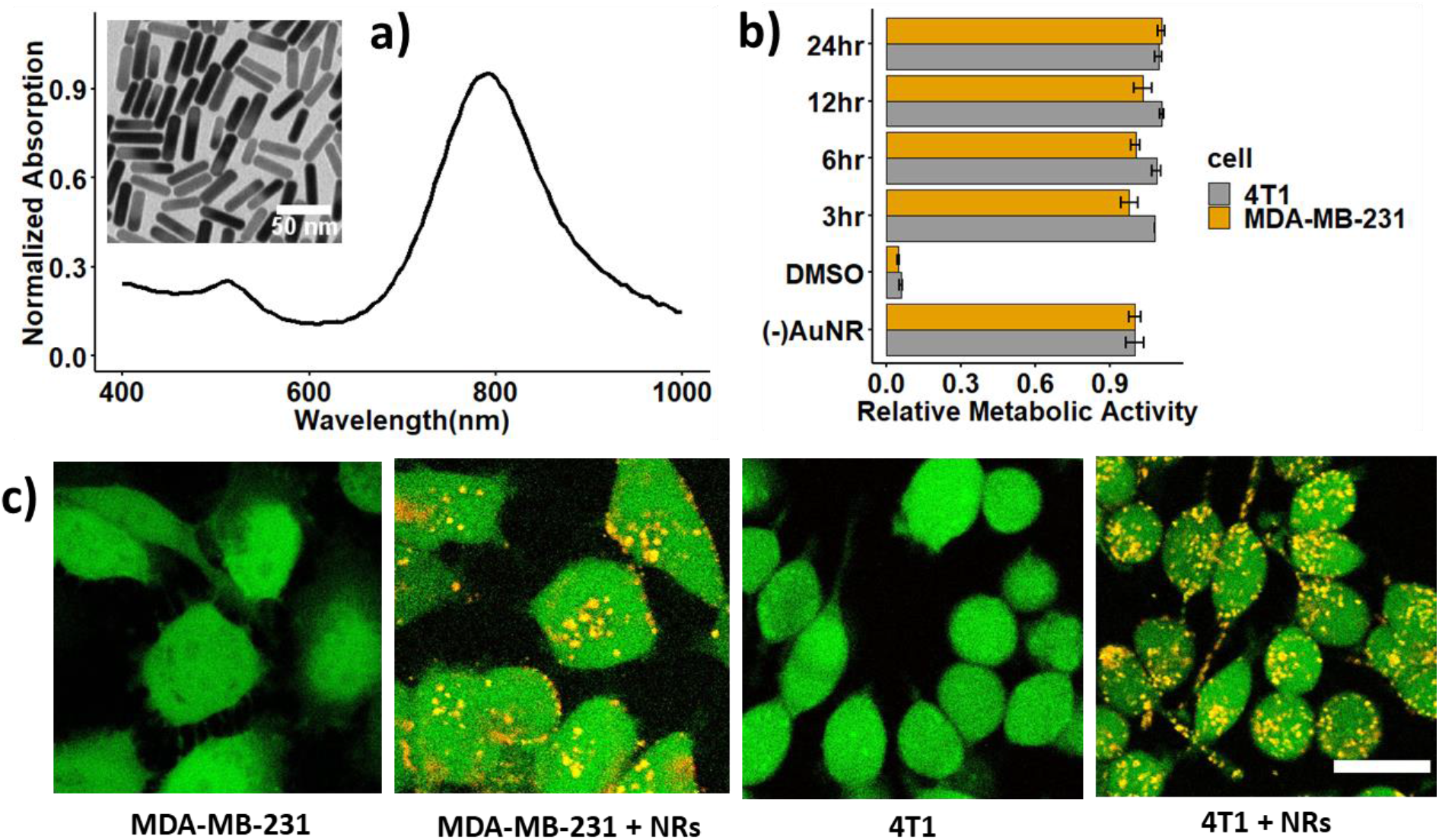
Gold nanorods (AuNRs) internalization in breast cancer cells: a) absorption spectrum of Mutab-coated AuNRs and TEM image of the nanorods (insert); b) relative metabolic activity of 4T1 and MDA-MB-231 cells incubated with AuNRs; c) two-photon images of MDA-MB-231 and 4T1 cells without and with AuNRs incubated for 6 hours. Scale bar: 20 μm

We determined the fluence threshold required for cell membrane disruption using calcein-AM and ethidium homodimer-1 (EthD-1) staining. The polyanionic calcein-AM can permeate through the membrane of live cells and produce an intense uniform green fluorescence in live cells. On the contrary, EthD-1 enters cells with damaged membranes, binds to nucleic acids, and provides a bright red fluorescence in dead cells. Figure 2a displays two-photon images of cells incubated with AuNRs after single-pulse nanosecond laser treatment at different fluences (0.7-5 J/cm^2^). We observed that the area of cell death occurs at the beam center at lower fluence and expands outwards from the beam center as laser fluence increases. The laser beam profile is approximately Gaussian (Figure S1); therefore, the ablation threshold is first exceeded in the beam center. The beam shape is not a perfect Gaussian and likely results in the observed irregularity in cell death areas. The fluence threshold required for membrane disruption is between 0.7 to 1.5 J/cm^2^, an order of magnitude higher than that reported in the literature^21^ of 0.07 J/cm^2^. This is likely a result of the 1064nm laser wavelength used here that operates off the resonance peak of the AuNRs at 788 nm. The damage threshold for cells without AuNRs is likely much higher, above 5 J/cm^2^, as we did not observe dead cells at this fluence (data not shown). We found a similar trend in 4T1 cells reported previously, where a fluence threshold for membrane disruption in cells with AuNRs is between 0.7 to 1.5 J/cm^2^, and the cell death area expands with increasing laser fluence^31^.

**Figure 2.**
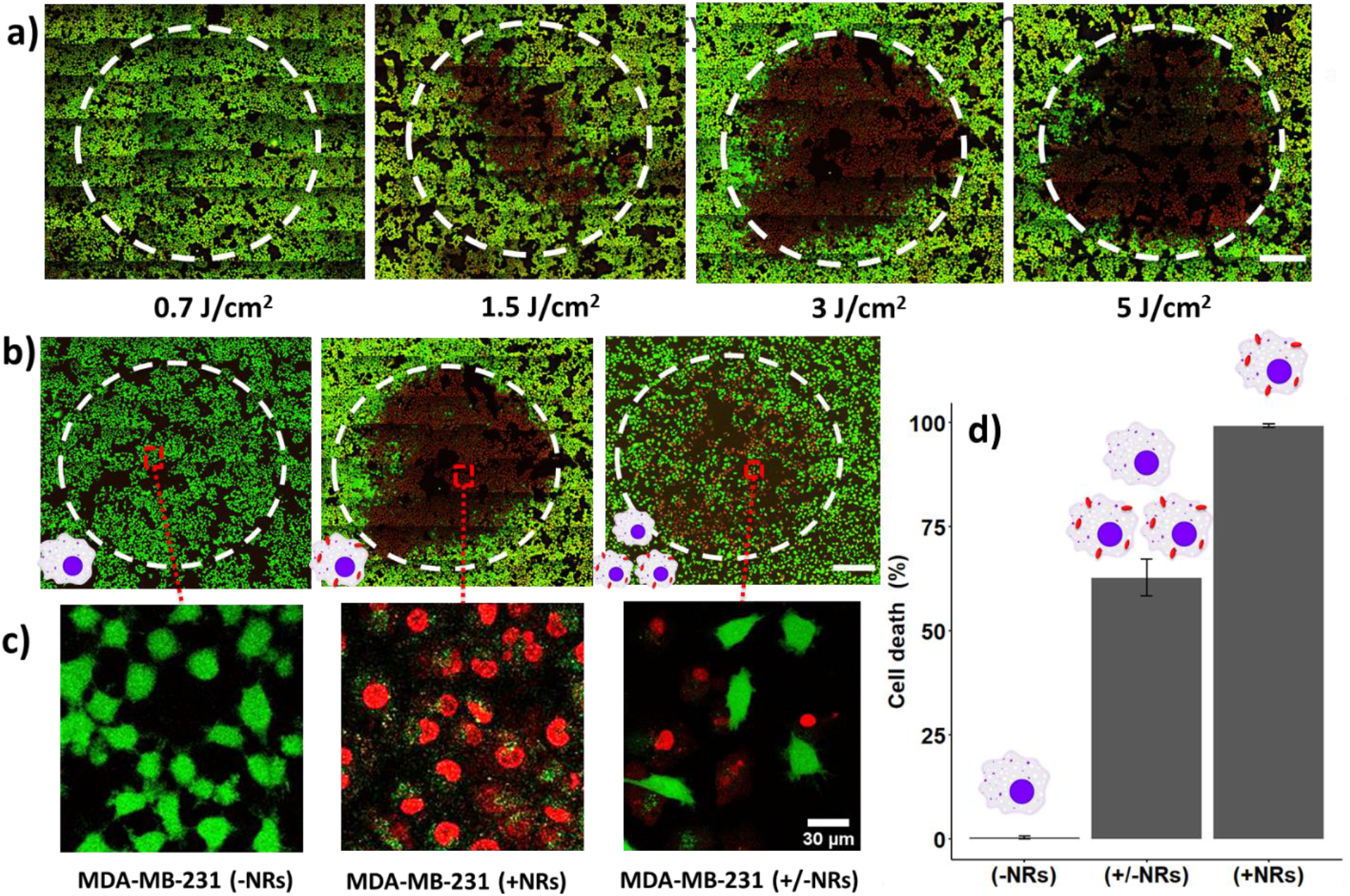
Cell death resulting from laser (1064 nm) irradiation of AuNRs-incubated MDA-MBA-231 cells: a) cell death with varying fluence from 0.7-5 J/cm^2^; b-d) cell death specific to irradiation of AuNRs-embedded MDA-MBA-231 cells at 3 J/cm^2^: without AuNRs incubation (left), with AuNRs incubation (middle) and with a mix of these two populations in 2:1 ratio (right). A two-photon microscope acquires the images with live cells stained in green (calcein-AM) and dead cells stained in red (ethidium homodimer-1). Beamwidth is highlighted with a white dashed line. Scale bar: 0.5 mm

To examine the specificity of laser treatment, we prepared three types of MDA-MB-231 cell populations: (a) without AuNRs, (b) with AuNRs, and (c) co-cultured cells with and without AuNRs at the ratio of 2:1. We irradiated cells at 3 J/cm^2^, followed by calcein-AM and EthD-1 staining (Figure 2b-c). We observed 0.3% of dead cells in the center of the beam for cells irradiated in the absence of AuNRs, 99% of dead cells for cells cultured with AuNRs, and 63% of dead cells for the group containing cells with and without AuNRs (Figure 2d). These percentages of cell death match well to the percentage of cells with AuNRs in the samples and demonstrate that within the laser beam, only AuNR-embedded cells were found dead while neighboring cells without AuNRs were intact. This observation is consistent with Pitsillides *et al*., who observed membrane disruption on cells with microparticles after nanosecond laser irradiation, while adjacent cells without microparticles were undamaged^22^. These results imply that this laser treatment is highly specific, affecting only the cells in direct contact with the AuNRs.

We visualized AuNRs-laser interaction in 4T1 cells using the custom inverted microscope setup shown in Figure 3a. The 1064 nm excitation laser beam was focused to a spot of 60 μm (full width half maximum) on the mono-layer 4T1 cells through the microscope objective (40x). The diffraction limit of the system is calculated at 0.25 μm. We used a high-speed camera (25,000 frames per second) to record videos of cells after single-pulse irradiation at 3 J/cm^2^ (supplemental video V1, V2; selected frames Figure 3b). The bubbles scatter light; hence, they appear as dark regions in image frames. We observed multiple bubbles expanding and collapsing on cells with AuNRs. Bubble diameters ranged between 0.8 and 3 μm (Figure 3b). No bubbles were observed after laser irradiation of cells without AuNRs (data not shown). Theoretical models and experimental measurements previously reported that nanosecond laser irradiation of single nanoparticle results in the generation of 0.1 - 0.5 μm bubbles^32–34^. The larger bubbles we observed may be due to AuNR clusters resulting from cell internalization. The bubble formation is evidence of the transient AuNRs absorption of the high energy laser pulse, which is then converted to mechanical disruption forces (high pressure and temperature) in cells. Moreover, these disruption forces are localized at the micrometer scale around AuNRs, which explains the high specificity of laser irradiation observed in Figure 2b-d.

**Figure 3.**
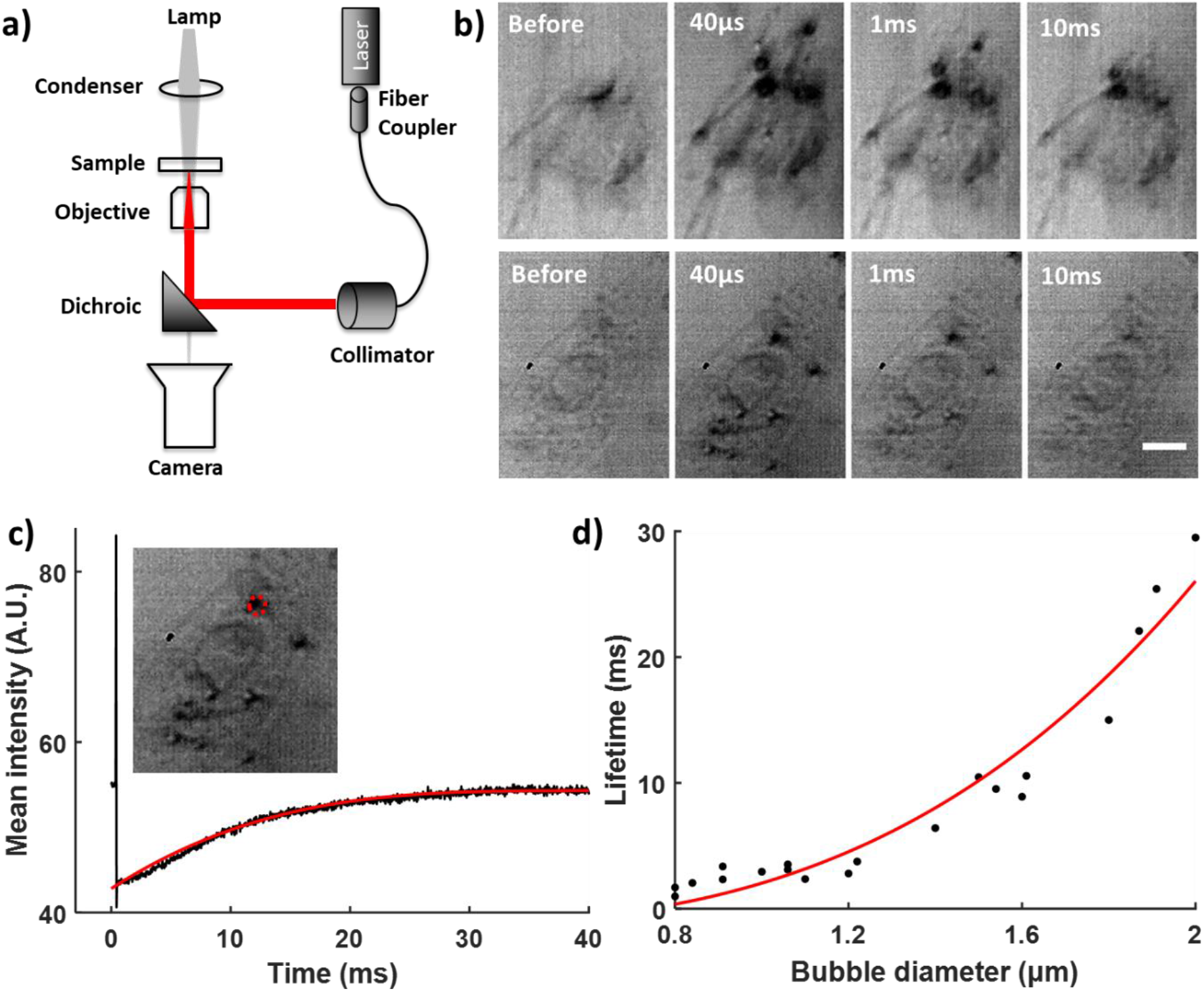
Imaging bubbles following rapid heating and water vaporization around AuNRs. (a) Assembly of the optical system to image bubbles, (b) montage of bubbles formed in two examples of 4T1 cells embedded with AuNRs after one pulse of laser irradiation at 3 J/cm^2^, (c) mean intensity of bubble over its time course with polynomial fitting (red line), the red dash line highlights the bubble pixels being monitored, (d) bubble’s lifetime vs. diameter with polynomial fitting (red line). Scale bar: 10 μm

We examined the bubble lifetime by monitoring the mean intensity of the bubble dark pixels over time (Figure 3c). At the time of laser irradiation, we observed a surge of intensity due to laser flash, followed by an instant sharp drop of intensity as the bubble formed. As the bubble collapsed, fewer dark pixels were present and the mean intensity recovered following a polynomial function as described in previous theoretical models studying gas bubble dissolution in liquid medium^35–38^. Zhang et al. derived the lifetime of a nanobubble τ as a function of its original radius *R*_0_ as follows^37^

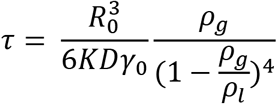

Where K is Henry’s law constant representing the gas solubility in liquid, D is the diffusion coefficient of the gas in the liquid, *γ*_0_ is the surface tension of liquid on a macroscopic scale, *ρ*_*g*_ and *ρ*_*l*_ are the density of gas and liquid, respectively. The lifetime of the bubbles recorded in cells (time that bubble intensity increases 1/e of its minimum-to-plateau difference) increased with the bubble’s diameter following a polynomial expression as described above (Figure 3d). We also observed that the bubble’s lifetimes are on the order of milliseconds, which are three orders of magnitude longer than the lifetime of similar-size bubbles in water (~300 ns) reported previously^32,34,39^. The long lifetime of bubbles in cells can be explained by the low gas solubility and diffusion coefficient in the cytoplasmic fluid. As cytoplasm fluid contains large biomolecules, its solubility of gases in it is expected to be much lower than that in water^40^. Furthermore, cytoplasmic diffusion of oxygen is two orders of magnitude lower than oxygen diffusion in water (~50 μm^2^/s in cytoplasmic fluid vs. 2500 μm^2^/s in water)^41^. While the inverted microscope images reveal the formation and collapse of bubbles in cells, we cannot confirm the onset of membrane disruption with the system. The cell membrane disruption observed in Figure 2 is likely the result of cavitation erosion, which occurs when bubbles collapse, generating re-entrant jet dynamics and emitting shock waves^42,43^, creating damage on the cell membrane.

To examine whether the nanobubbles can trigger immunogenic cell death, we characterized the release of damage-associated patterns (DAMPs) including heat shock protein 70 (HSP70), chaperone protein human high mobility group box 1 (HMGB1) and adenosine triphosphate (ATP) in the extracellular environment by ELISA and bioluminescence assays (Figure 4). The MDA-MB-231 and 4T1 cells were treated with Doxorubicin 1μg/ml and 10μg/ml, respectively, for 24 hours as a positive control. We observed increased release of all three types of DAMPs in both cell lines in groups with AuNRs and laser irradiation. The 4T1 cells released more DAMPs than the MDA-MB-231 due to its higher metabolic activity. We also observed two trends of DAMPs secretion: the release of HSP70 and HMGB1 increased with time while the ATP quenched quickly with time (Figure 4b). After membrane disruption from laser irradiation, the DAMPs proteins were released gradually into the extracellular environment as cells follow the necrosis pathway. On the contrary, most ATP molecules inside the cells were secreted immediately after laser irradiation. These ATP molecules were unstable in the extracellular environment and hence, were quickly lessened after one hour. Laser irradiation triggered ATP release instantly, creating a surge of ATP in the extracellular environment in contrast with doxorubicin treatment, where ATP was released gradually. As a result, we observed a much higher amount of ATP in the laser-treated group than the doxorubicin-treated group at the time of measurement.

**Figure 4.**
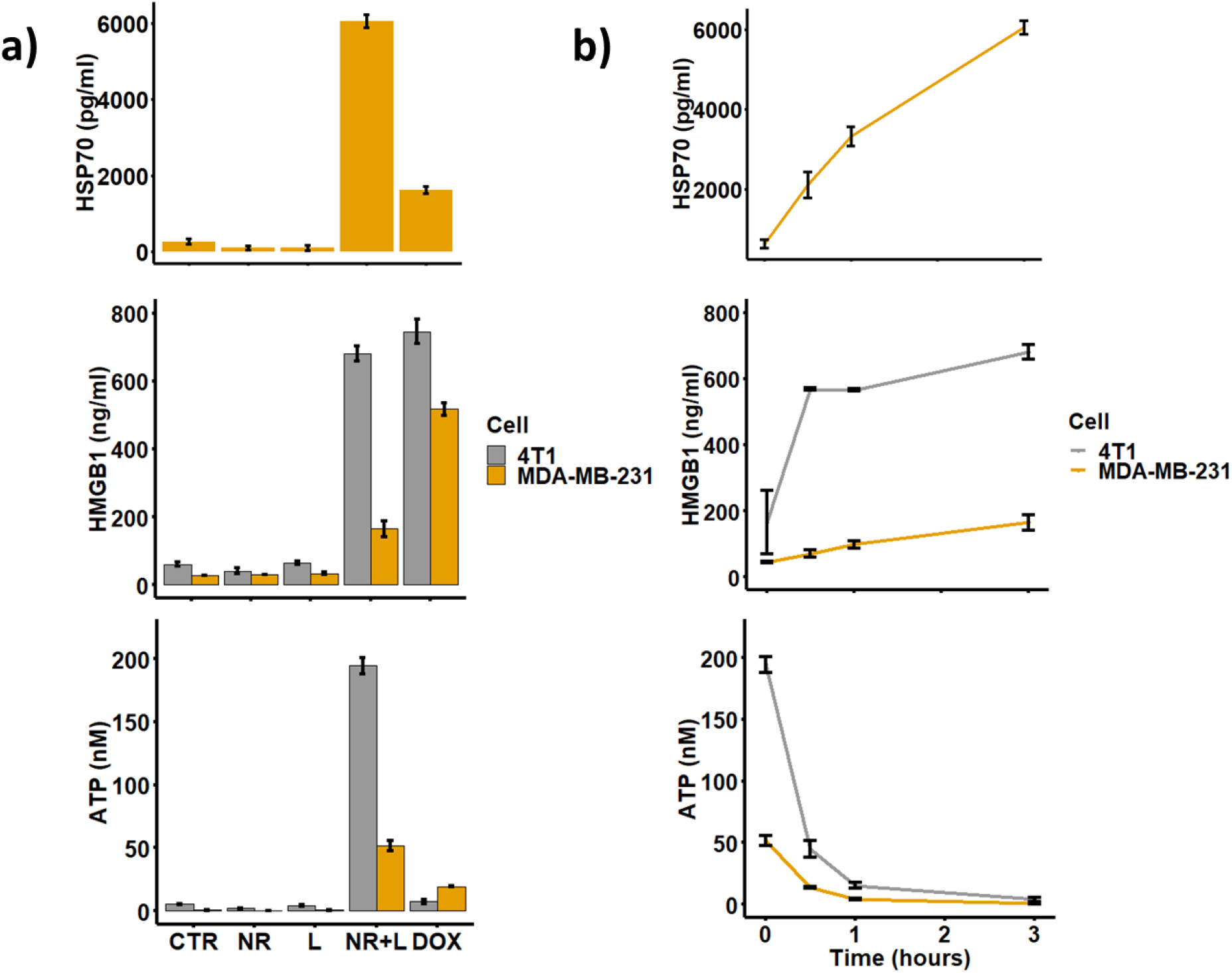
Extracellular release of damage associated patterns from laser irradiation of MDA-MB-231 and 4T1 cells at 3 J/cm^2^ and 2 cm^2^ of cells area per sample (50% of the well area), a) HSP70, HMGB1, and ATP; b) Time-dependent release of DAMPs after laser irradiation. Groups: CTR: cells without any treatment, NR: cells incubated with AuNRs, L: cells irradiated with a laser, NR+L: cell incubated with AuNRs and irradiated with a laser, DOX: cell treated with doxorubicin for 24 hours as the positive control. The number of samples per group n = 3.

To verify whether dendritic cells are activated with the presence of DAMPs signals released from laser irradiation, we co-cultured 4T1 cells and dendritic cells (DCs) in a trans-well system (Figure 5a). Bone marrow-derived dendritic cells (DCs) from BALB/c mice housed were cultured as per the Lutz method, with the addition of IL-4.^44–46^ A significant increase in the percentage of mature DCs (Cd11c+ MHCII+ and CD86+) was observed in a group of laser irradiation of AuNRs-embedded 4T1 cells (Figure 5b, c). This increase is mostly driven by an upregulation in major histocompatibility complex (MHCII), which increases several folds during dendritic cell maturation^47^. We did not observe an increase in the percentage of mature dendritic cells when the 4T1 cells with AuNRs were irradiated for one time. We believe the ratio of 4T1 cells over DCs is crucial for activating DCs from DAMPs. This ratio is very skewed towards tumor cells *in vivo* as DCs are rare population in tissue (<1%)^48^, while our simulated experiment is oppositely skewed due to the design of the transwell system (one 4T1 cell per DC). By irradiating 4T1 cells incubated with AuNRs more than once, more DAMPs per DCs are attained, thus increase the chance of activating DCs. We expect a more favorable outcome *in vivo* experiments because the ratio of DAMPs per DC will be significantly higher.

**Figure 5.**
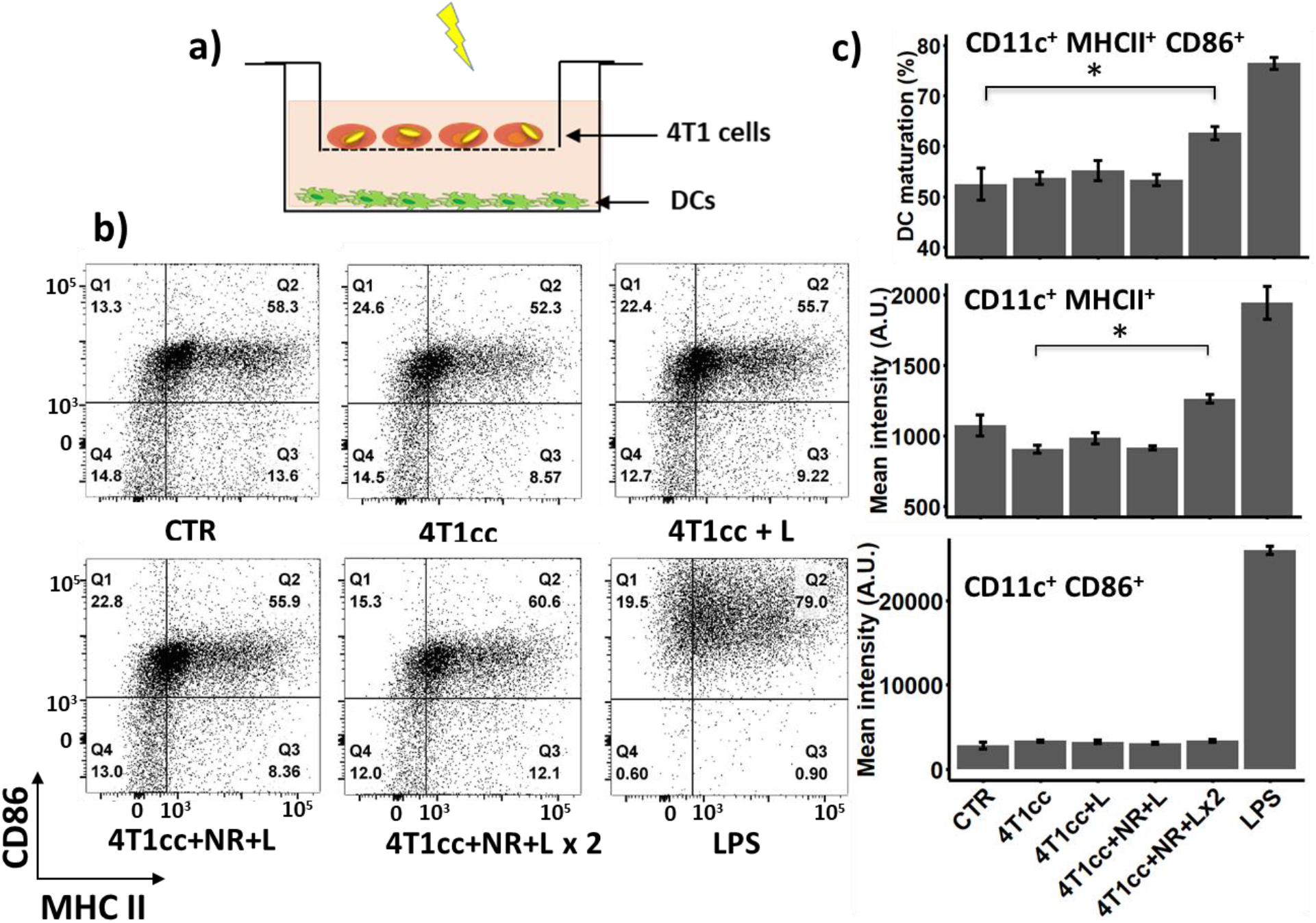
Activation of dendritic cells from irradiation of AuNRs-embedded 4T1 cells a) experimental layout describing 4T1 co-cultured with DCs in a transwell setting, b) Dot plot of DCs expressing MHC II and CD86, c) Percentage of mature dendritic cells as CD11c+ MHCII+ and CD86+, and median intensity of DCs that express MHCII and CD86. Six groups of dendritic cells: CTR: DCs without treatment; 4T1cc: DCs co-cultured with 4T1 cells; 4T1cc+L: DCs cultured with irradiated 4T1 cells; 4T1cc+NR+L: DCs co-cultured with irradiated AuNRs-embedded 4T1 cells; 4T1cc+NR+L x 2: DCs co-cultured with twice irradiated AuNRs-embedded 4T1 cells, the second irradiation is 12 hours after the first irradiation; LPS: DCs treated with LPS at 1μg/ml for 12 hours. Number of samples per group n = 3. Statistical analysis was performed with analysis variance (ANOVA) in combination with Tukey test, * means p-value < 0.05

In conclusion, we observed a single 1064nm nanosecond laser pulse in combination with gold nanorods eradicates breast cancer cells and induces immunogenic cell death. We detected cell death from membrane disruption after laser irradiation, which only happened in cells with nanorods, while neighboring cells without nanorods were left intact. We also observed bubbles and discrete cellular damage around the bubbles as the result of nanorod-laser interaction. We demonstrated that DAMPs released in the extracellular environment are enhanced in the laser treatment group. With the presence of DAMPs released from laser irradiation, dendritic cell activation was also increased. Overall, we determined that nanosecond pulsed laser irradiation provided a fast and highly specific approach to eradicate tumor cells and induce markers of immunogenic cell death. This study provides supporting experimental evidence of the concept that laser nanobubbles trigger immunogenic cell death in cancer cells and is a candidate approach as a combination strategy for immunotherapy.

## ASSOCIATED CONTENT

### Supporting Information

Details about the laser beam profile, calreticulin redistribution from nanobubbles, dendritic cell maturation induced from the supernatant of treated groups, and gating strategy for Figure 5.

## ACKNOWLEDGMENTS

The work was supported by the Cancer Prevention and Research Institute of Texas (CPRIT RP130702), Texas Health Catalyst, and Texas 4000. The authors gratefully acknowledge funding through the American Heart Association (17IRG33410888), the DOD CDMRP (W81XWH-16-1-0580; W81XWH-16-1-0582) and the National Institutes of Health (1R21EB023551-01; 1R21EB024147-01A1; 1R01HL141761-01) to ABB. The authors sincerely appreciate the fruitful discussion with professor Preston Wilson and Dr. Emil Sobol on the topic of nanobubble dynamics and discussion with professor Rongze Lu on the subject of dendritic cell activation. We would like to thank Dr. Adrian Spencer and Daniel Chavarria from Baker lab for help with the MDA-MB-231 cell line; Scott Jenney and Dr. Bharadwaj Muralidharan from Milner lab for assistance with the optical fibers and alignment; Dr. Kabir Dhada from Suggs lab for support with the Nd: YAG 1064 nm laser.

---

**Figure S1.**
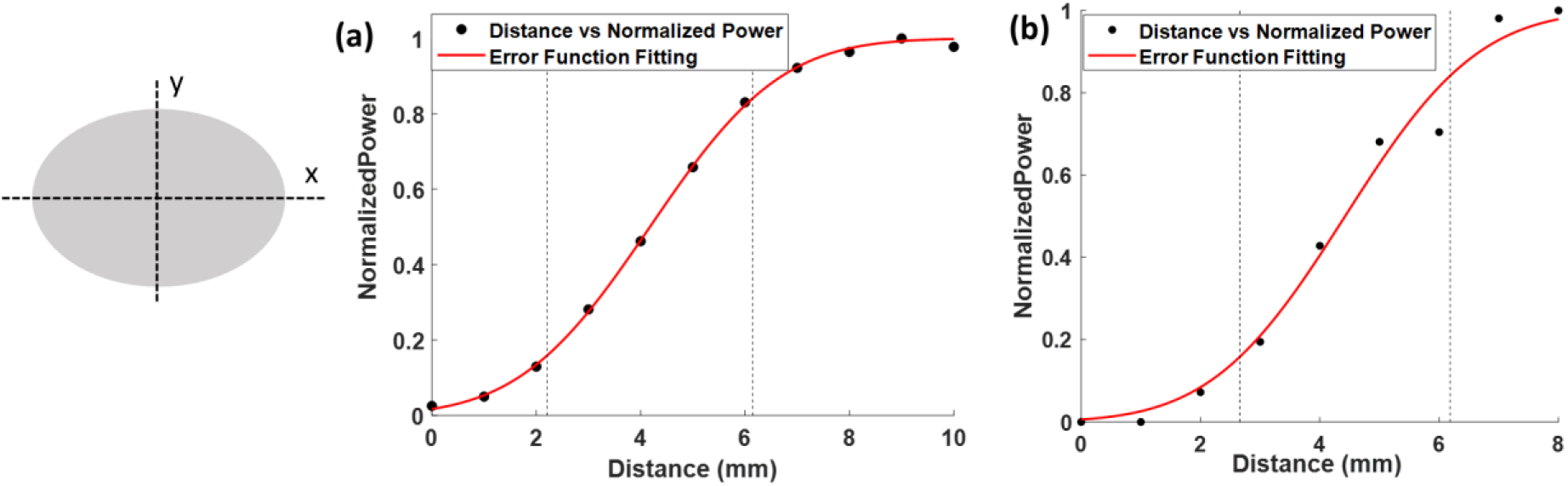
Laser beam profiling with knife-edge method in the x-direction (a) and y-direction (b). Briefly, we moved a knife blade along either x or y direction and recorded the power of the laser beam that was not covered by the knife blade. These recorded values formed the cumulative distribution function of a Gaussian beam and hence were fitted to an error function. The beam waist (full width half maximum) was labeled between the two vertical dashed lines.

**Figure S2.**
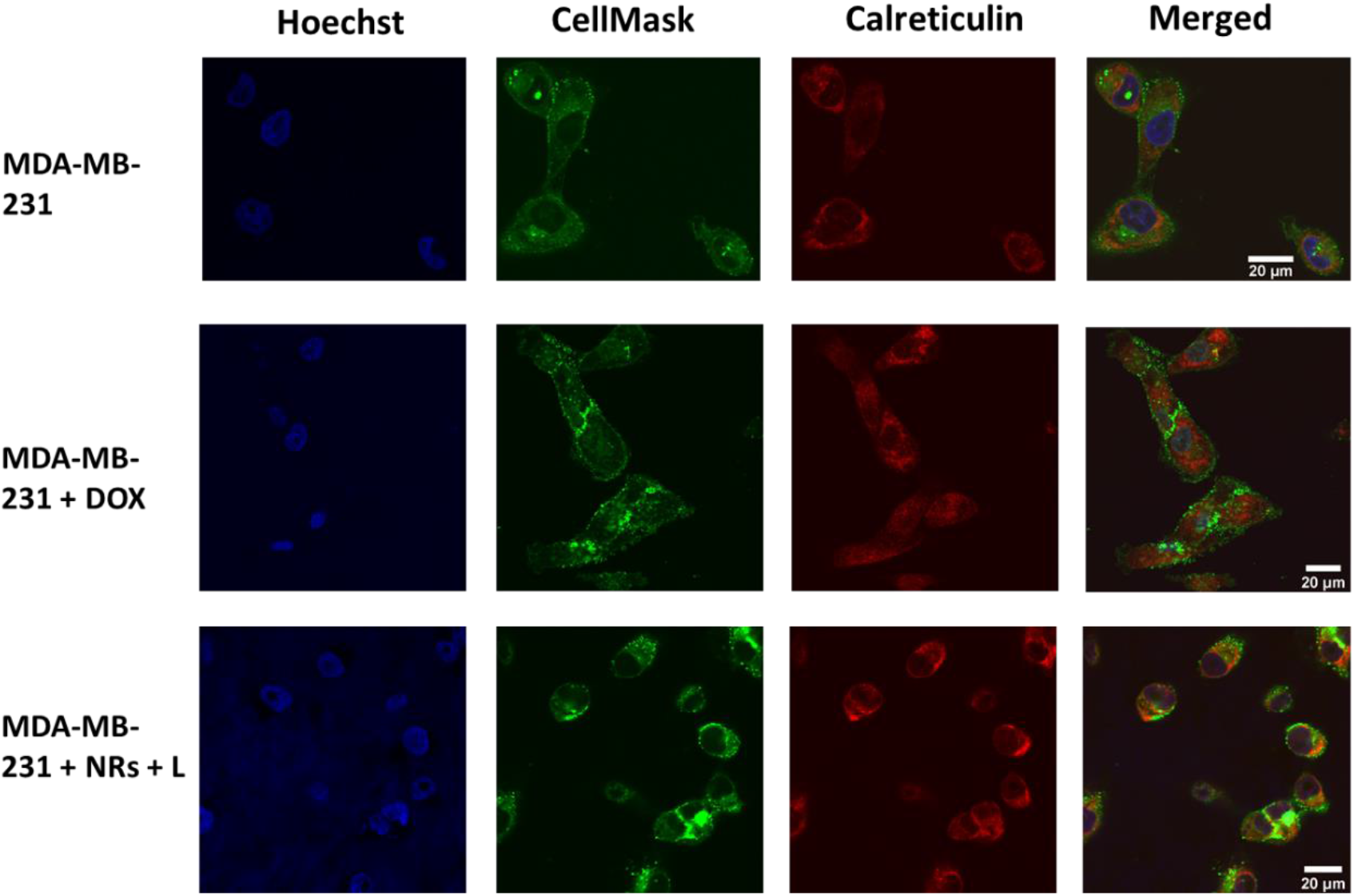
Characterization of calreticulin relocation with confocal imaging. We labeled cell nucleus with Hoechst (blue), cell membrane with Cellmask (green), and calreticulin with AF647 conjugated antibodies (red). We observed calreticulin colocalizing with membrane label after laser irradiation of MDA-MB-231 cells with AuNRs.

**Figure S3.**
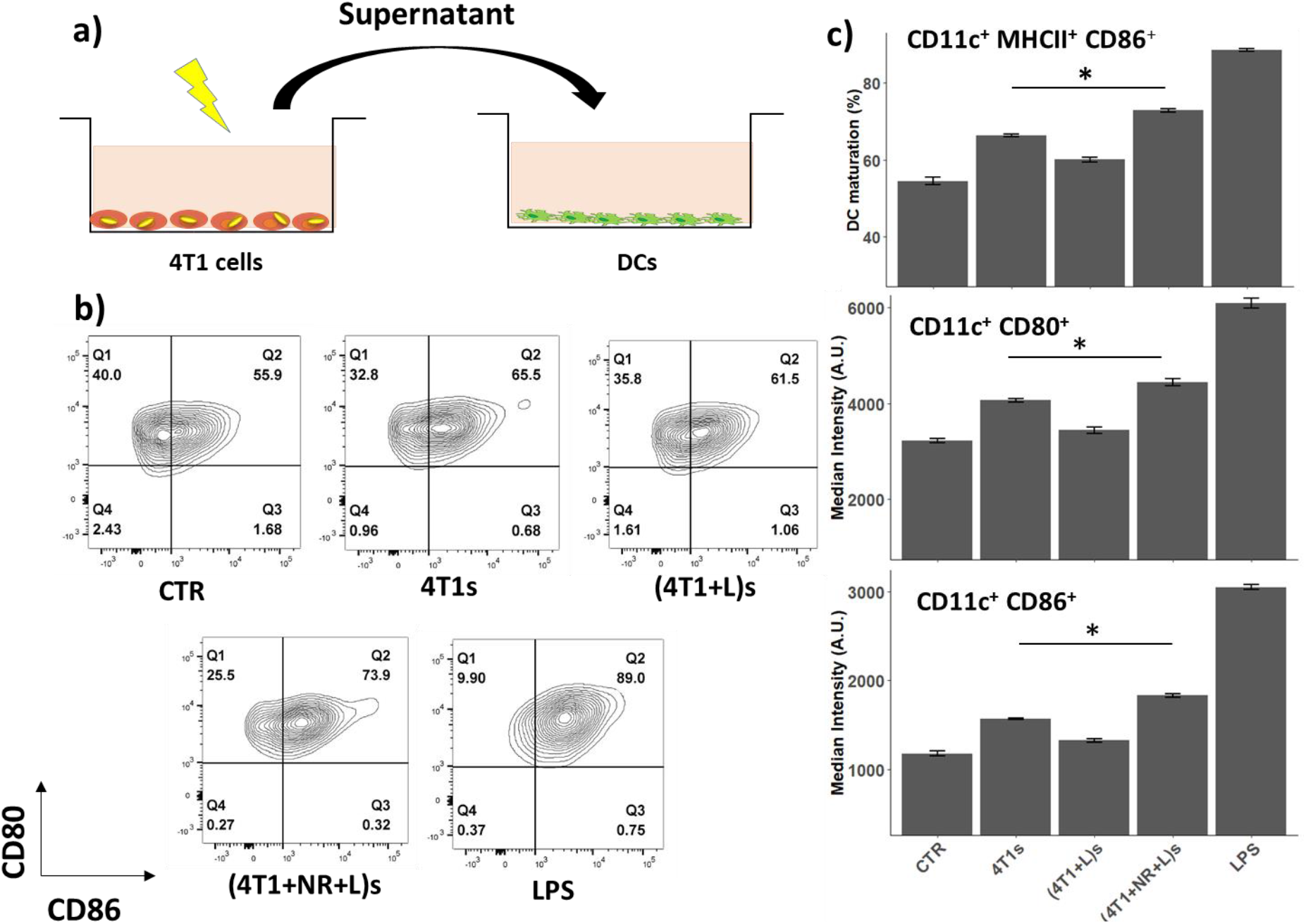
Activation of dendritic cells from irradiation of AuNRs-embedded 4T1 cells a) experimental layout describing 4T1 supernatant addition to dendritic cells (DCs) b) contour plot of DCs expressing CD80 and CD86, c) Percentage of mature dendritic cells as CD11c+ CD80+ and CD86+, and median intensity of DCs that express CD80 and CD86. Five groups of dendritic cells: CTR: DCs without treatment, 4T1s: DCs treated with supernatant from 4T1 cells, (4T1+L)s: DCs treated with supernatant from irradiated 4T1 cells, (4T1+NR+L)s: DCs treated with supernatant from irradiated AuNRs-embedded 4T1 cells, LPS: DCs treated with LPS. Number of samples per group n = 3. * means p-value < 0.05

**Figure S4.**
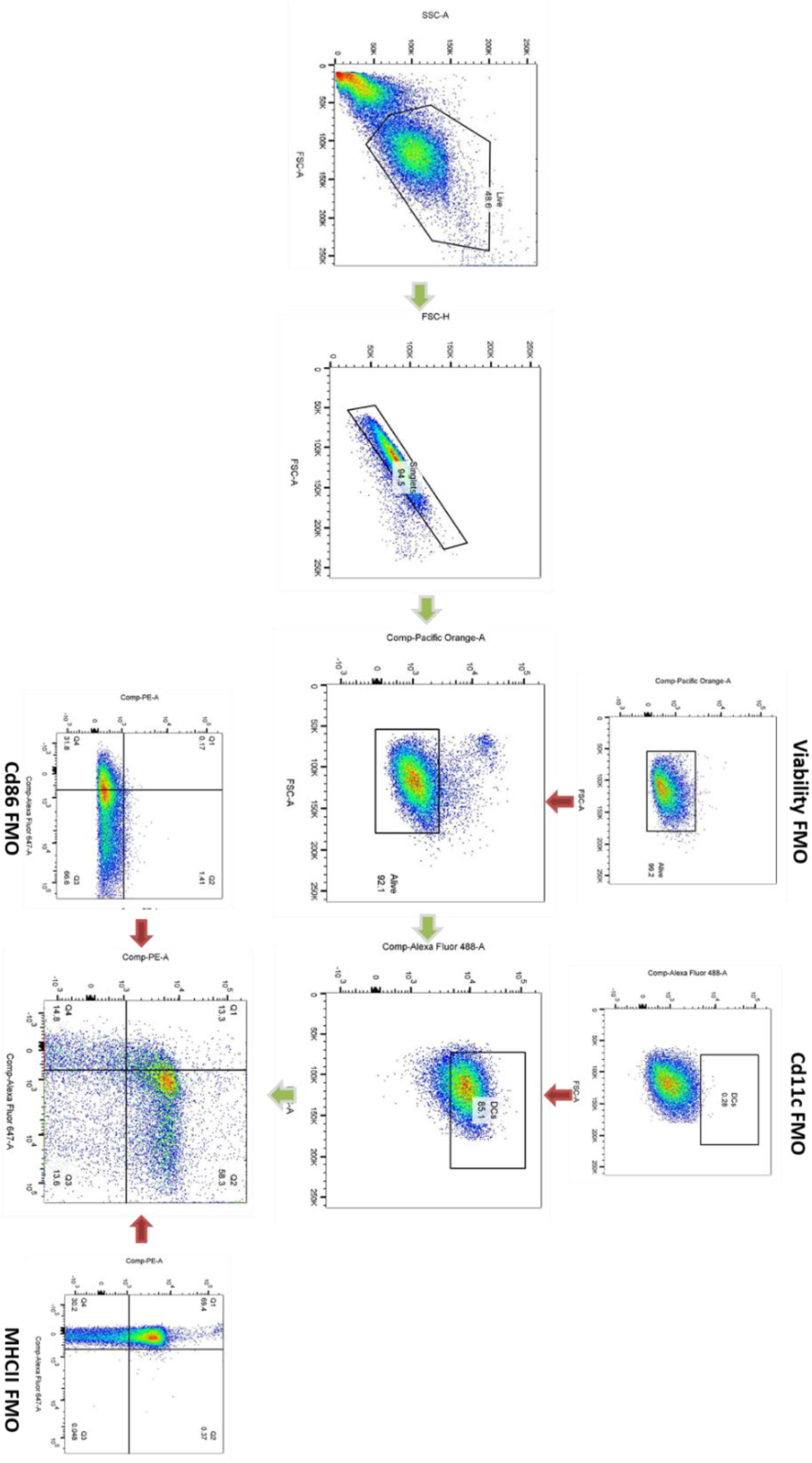
Gating strategy for dendritic cell maturation. We first gated out the debris following by gating the singlets. We used a viability dye to gate live cells. Cd11c is the markers for dendritic cells, while Cd86 and MHCII are the markers for mature dendritic cells. All gates with fluorophores were defined by fluorescence minus one (FMO) samples.

